# Single-Cell Resolution DESI Mass Spectrometry Imaging through 10-Fold Sample Expansion

**DOI:** 10.1101/2024.10.21.619369

**Authors:** Chengyi Xie, Jianing Wang, Xin Diao, Xiaoxiao Wang, Zongwei Cai

**Affiliations:** State Key Laboratory of Environmental and Biological Analysis, Department of Chemistry, Hong Kong Baptist University, Hong Kong SAR, China; College of Science, Eastern Institute of Technology, Ningbo, Zhejiang, China

## Abstract

Desorption electrospray ionization mass spectrometry imaging (DESI MSI) is a valuable tool for label-free, spatially resolved molecular analysis of biological tissues. However, its spatial resolution has been limited to tens of micrometers due to constraints in spray geometry and solvent flow, hindering single-cell imaging. Inspired by expansion microscopy, we developed a tenfold DESI expansion MSI (10x-DESI ExMSI) technique to achieve cellular-level resolution. In this study, tissue sections were embedded in a swellable polyelectrolyte gel, and physically enlarged by approximately tenfold without compromising molecular integrity. After drying of the sample, mass spectrometry imaging was performed using a custom-built DESI ion source coupled with an Orbitrap mass spectrometer. Using mouse brain tissues as representative samples, 10x-DESI ExMSI achieved a spatial resolution of approximately 5 µm, comparable to the highest levels attained by advanced commercial MALDI-MSI systems. Enhanced resolution allowed clear visualization of fine structural details, such as the granule cell layer of the dentate gyrus and the pyramidal layer of the Cornu Ammonis, which were unresolved in unexpanded samples under a 100 μm step size. Lipid profiling demonstrated high retention and detection efficiency, confirming the technique’s suitability for comprehensive molecular imaging. The 10x-DESI ExMSI significantly advances DESI MSI by enabling single-cell level resolution without modifying existing mass spectrometers. The method is simple, cost-effective, and easily implementable in most laboratories, making high-resolution DESI MSI accessible for a wide range of biological applications. This approach overcomes longstanding limitations of DESI imaging and holds great potential for detailed molecular studies at the cellular level.

## Introduction

The progression of modern biology relies on the development of advanced analytical technologies that offer detailed molecular distribution and structural information within tissues and cells. Mass spectrometry imaging (MSI) has become a critical tool, offering label-free, spatially resolved molecular analysis across diverse biological and chemical systems. MSI enables the simultaneous detection of thousands of molecular ions, making it particularly valuable in biology, medical sciences, and drug discovery^1,2^. MSI techniques can be broadly classified into two categories: molecular MSI, which uses soft ionization to preserve intact molecular ions, and elemental or atomic MSI, which employs hard ionization methods. While atomic MSI techniques such as secondary ion mass spectrometry (SIMS) and laser desorption ionization (LDI) achieve high spatial resolution using high-energy ion beams or lasers, they primarily produce elemental ions or small molecular fragments^3^. This extensive fragmentation limits their utility in biological tissues, where preserving intact biomolecules is essential for comprehensive molecular profiling. Despite their ability to achieve sub-micron resolution, these techniques are less effective in revealing the rich molecular information found in biological samples. Therefore, developing high-resolution molecular MSI techniques is crucial for advancing biological imaging, as they provide the molecular detail necessary for understanding complex biological systems.

Molecular MSI techniques, including matrix-assisted laser desorption/ionization (MALDI) and desorption electrospray ionization (DESI), rely on soft ionization processes that retain intact biomolecules, making them the primary methods for biological tissue imaging. MALDI MSI, one of the most widely adopted MSI techniques, can achieve spatial resolutions up to 5 μm in commercial systems, enabling high-resolution imaging of tissue sections at single-cell resolution. DESI, introduced by Cooks and colleagues in 2004^4^, offers a complementary approach as an ambient ionization technique that requires minimal sample preparation. In DESI, charged droplets are pneumatically accelerated to the sample surface, where subsequently splashed secondary droplets carrying desorbed and ionized analytes, which are then transferred to the mass spectrometer for analysis. The simplicity of the DESI setup and its ability to perform in-situ analyses without matrix application have led to widespread adoption in diverse research fields^5–7^. However, DESI MSI typically achieves lower spatial resolution than MALDI MSI, with effective spatial resolution often exceeding 100 μm. Efforts to enhance DESI’s spatial resolution, such as optimizing experimental parameters, have reduced the pixel size to 25-30 μm in some cases^8^. Nevertheless, the inherent spray geometry and continuous nature of the solvent flow limit the development of high-resolution DESI MSI, restricting its application to cellular-level molecular imaging. To address the limitations of DESI’s spatial resolution, nanoDESI, a modified version of DESI, has been developed^9^. NanoDESI utilizes a small liquid bridge between the sample and the mass spectrometer, effectively avoiding the detrimental effects of the electrospray cone angle, thus enabling better spatial resolution. NanoDESI MSI has achieved resolution as low as 10 μm^10^, representing a notable improvement over conventional DESI MSI. However, nanoDESI MSI faces significant operational challenges, particularly in achieving stable and continuous long-term high-resolution operation. For example, at acquisition resolutions around 10 μm, nanoDESI MSI often experiences issues such as filament-like lateral oversampling and signal interruptions, which severely impact data quality and consistency. DESI MSI, with modifications to the sprayer design, could reduce the spatial resolution to around 20 μm^11^. The inherent complexity of maintaining droplet stability and controlling the solvent plume at these fine scales presents persistent challenges.

For more reliable and stable acquisition, robust and easily operable DESI MSI systems typically achieve an effective spatial resolution above 40 μm under general conditions. In contrast, MALDI imaging routinely achieves stable and reproducible spatial resolutions at 10 μm. Additionally, some of the most advanced MALDI imaging platforms can achieve spatial resolutions as fine as 5 μm, further enhancing image detail and accuracy. This emphasizes a significant performance gap between DESI and MALDI imaging when targeting cellular-level resolution, with MALDI providing more reliable, high-resolution imaging and greater stability in practical applications.

Inspired by expansion microscopy^12^, which enables high-resolution imaging by physically enlarging biological samples, we have pioneered the expansion mass spectrometry imaging (ExMSI) strategy to achieve ultra-high spatial resolution^13^. In this study, we introduce a 10x DESI ExMSI system, which integrates a regular custom-built DESI ion source with a newly developed workflow. By implementing a 10-fold expansion factor, we achieve single-cell level resolution in DESI MSI, enhancing its spatial resolution to approximately 5 μm-comparable to the leading commercial MALDI systems. The system is simple, cost-effective, and easily implemented in most labsoratories without modifying existing mass spectrometers. This expansion mass spectrometry strategy overcomes long-standing DESI imaging limitations, representing a significant advancement that makes high-resolution DESI MSI accessible and reliable.

### Results and Discussion

#### Expansion-DESI MSI

The limited spatial resolution of conventional DESI MSI has restricted its application in discerning fine structure details within biological tissues. To address this challenge, we have developed a novel tenfold magnification expansion-DESI MSI (10x DESI ExMSI) workflow, significantly enhancing the effective spatial resolution of DESI MSI (Fig. 1a). Briefly, a 50 μm thick fixed tissue slide was prepared by cryosectioning. The tissue section was placed in a custom-built slide chamber and embedded in a swellable hydrogel. The chamber was incubated at 37 ^°^C in a humidified environment to complete the gelation process. During gelation, proteins were anchored to the gel network under well-controlled conditions. After gelation, the sample was fully expanded in water until no further expansion occurred. The tissue was then fully dried. Before DESI imaging, an optional step is the application of solvent onto the sample surface, which is beneficial for the pre-extraction of analytes. To determine the effective expansion factor, the lengths were measured before expansion and after drying (Supplementary Fig. 1). As shown in the figure, the bottom of the mouse hippocampus slide was measured 5.4 mm before expansion, which became 51mm after expansion and drying, indicating a 9.4-fold expansion. Finally, MS imaging was conducted using our cost-effective homebuilt DESI source, coupled with an Orbitrap mass spectrometer. The acquired MSI data were analyzed and visualized by SCILS Lab MVS.

**Fig. 1:**
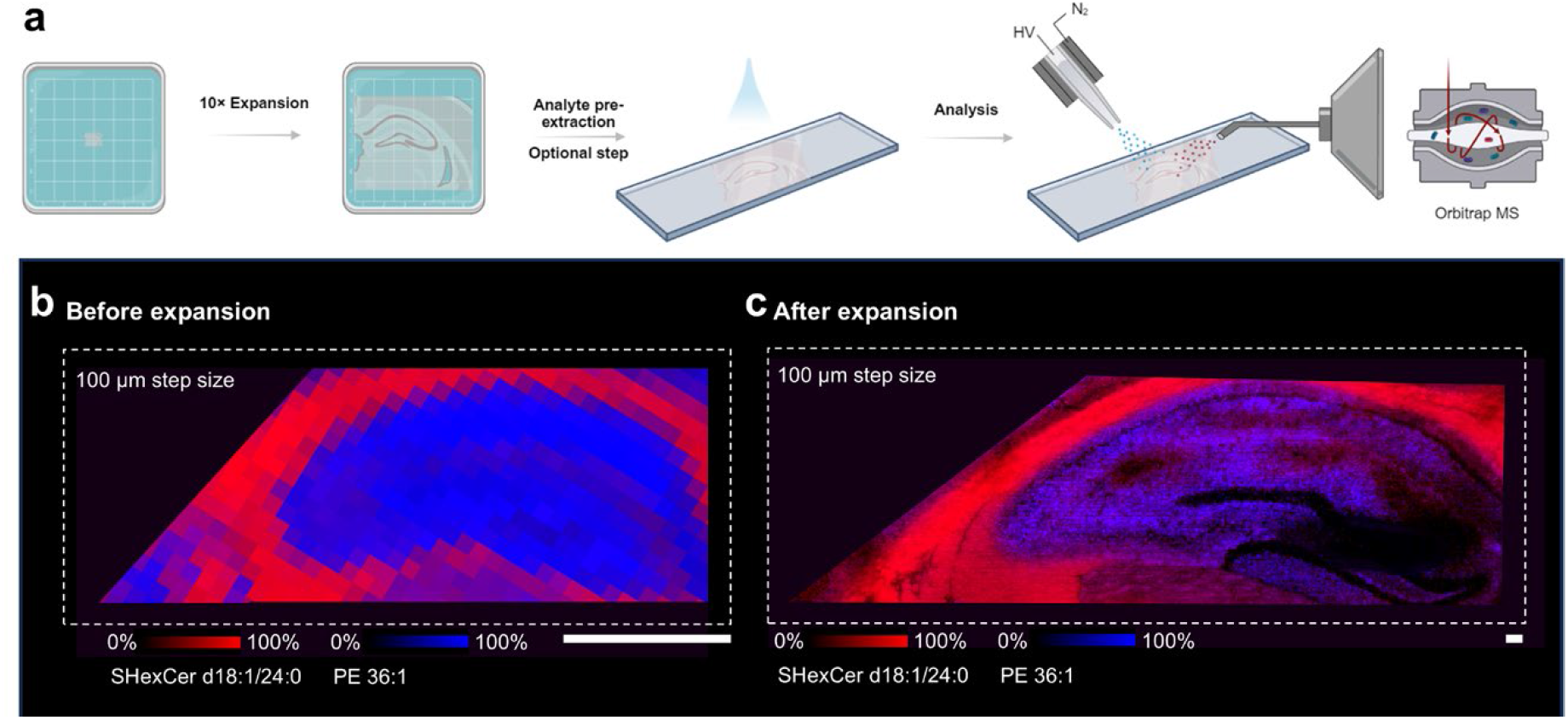
Schematic Overview of expansion DESI MSI. a) workflow of ten-fold magnification expansion DESI MSI includes tissue expansion, analyte pre-extraction(optional step), and DESI MSI analysis. b and c) overlay ion images of SHexCer d18:1/24:0 (m/z 888.62) and PE 36:1 (m/z 744.56) obtained for expansion mouse hippocampus and conventional counterpart by DESI MSI. Scale bar represents 1mm.

Mouse hippocampus was used as a representative example to demonstrate the effectiveness of expansion DESI imaging with a vertical step size set to 100 μm. The overlay ion images of ions [SHexCer d18:1/24:0 – H]^-^ and [PE 36:1 – H]^-^ at m/z 888.63 and 744.55 for expanded mouse hippocampus and the unexpanded counterpart are shown in Fig. 1b. Notably, expansion DESI MSI provides much clearer structural visualization compared to conventional DESI MSI under the same vertical step size. Specifically, while the granule cell layer of dentate gyrus (DG) is clearly resolved after expansion, the corresponding structure in the normal sample appears ambiguous. Additionally, the pyramidal layer of Cornu Ammonis (CA), which is unresolved by conventional DESI MSI, is clearly differentiated after expansion. The result confirm that our expansion strategy leads to a significant improvement in spatial resolution for DESI MSI, enabling the visualization of fine structural details previously unattainable with standard DESI techniques.

Smaller step sizes and slower imaging scan rates will lead to a higher spatial resolution for DESI MSI^8^. We first assessed the spatial resolution achievable by our homebuilt DESI platform using a normal mouse hippocampus. In DESI MSI, the attainable spatial resolution can be determined by measuring the distance over which the most defined features in the image transition from 20% to 80% in relative signal intensity^10,14^. As shown in Supplementary Fig. 2c, the distance is measured at 36 μm from the relative abundance of 20% to 80% of the maximum ion intensity when scanning a single line. An averaged spatial resolution of 48 μm could be achieved using a step size of 40 μm at a lateral speed of 19.2 μm/s (Supplementary Fig. 2d). To further investigate the impact of smaller step size on the effective spatial resolution, the step size was set to 25 μm. Although oversampling occurred, the average spatial resolution was decreased to 39 μm (Supplementary Fig. 2d). In the overlay ion images of [PS 40:6 – H]^-^, [PI 38:4 – H]^-^, and [SHexCer d18:1/24:0 – H]^-^, the 48 μm spatial resolution provides the ability to resolve the granule cell layer in DG region and the pyramidal layer in CA region (Supplementary Fig. 2e). The 39 μm spatial resolution provides clearer structure visualization of mouse hippocampus. However, the reduced step size significantly increased the acquisition time, making it less practical for routine analysis. To maintain a balance between spatial resolution and acquisition time, we recommend a 40 μm step size for DESI imaging experiments, with the option to use a 25 μm step size when necessary.

#### High spatial resolution Ex-DESI MSI

As the proof of achieving higher spatial resolution using smaller step size and lower scanning speed (Supplementary Fig. 2e) and the improvement of spatial resolution after expansion (Fig. 1b and Fig. 1c), we further investigate the ability of Ex-DESI to resolve detailed structure under higher spatial resolution. A scanning settings of a line speed of 19.2 μm/s and a step size of 40 μm, whose spatial resolution was calculated to about 48 μm (Fig. 1d), were used for the visualization of the striatum region in mouse brain. After a near 10-fold expansion, an equivalent spatial resolution of near 5 μm can be achieved under the scanning settings, approaching the upper limitation of spatial resolution for commercial MALDI instrument (Fig. 2c). For comparison, a conventional (unexpanded) sample was acquired at the same instrumental settings (Fig. 2a), and another expansion sample was measured using a scanning line speed of 76.9 μm/s and a step size of 100 μm (Fig. 2b). As shown in Fig. 2a, the 48 μm spatial resolution is unable to resolve the detailed structure of striatum dorsal region presented in the ion image of [SHexCer d18:1/24:0 – H]^+^ at m/z 888.62. On the contrary, the expansion DESI MSI readily resolved the fibrous structures even with a larger step size of 100 μm (Fig. 2b). The fibrous structure is visualized more clearly by an equivalent spatial resolution of about 5 μm (Fig. 2c) when further reduce the step size to 40 μm. The ion image measured by expansion DESI MSI can be aligned with immunofluorescence (IF) staining of myelin basic protein (MBP) (Fig. 2d), which highlights the distribution of myelin in the nervous system.

**Fig. 2:**
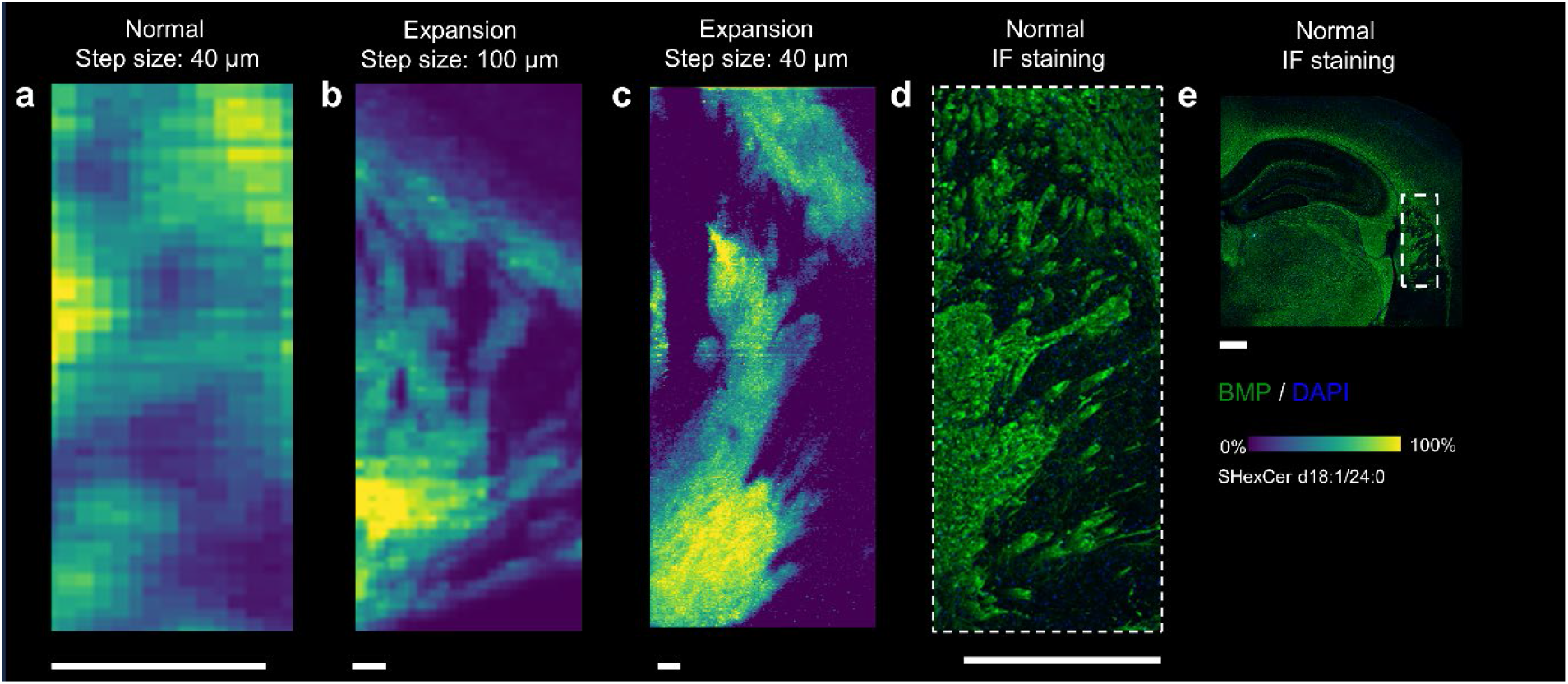
High lateral resolution expansion DESI MSI of mouse striatum. Ion image of [SHexCer d18:1/24:0 – H]^-^ at m/z 888.62 acquired by a) conventional DESI MSI at a step size of 40 μm and line speed of 19.2 μm/s, b) expansion DESI MSI at a step size of 100 μm and scanning line speed of 76.9 μm/s, and c) expansion DESI MSI at a step size of 40 μm and line speed of 19.2 μm/s. e) optical image of mouse brain at coronal plane after staining using MBP and DAPI. d) expanded view of the region indicated with a white box in e. Scale bar represents 500 μm.

To further demonstrate the effectiveness of high spatial resolution Ex-DESI MSI, we analyzed both expanded and normal mouse cerebellums using a step size of 40 μm and line speed of 19.2 μm/s. In the expanded sample (Fig. 3c), the white matter layer, granule layer,and molecular layer were fully resolved in the overlay ion images of [SHexCer d18:1/24:0 − H]^−^ and [PI 38:4 − H]^−^ at m/z 888.62 and 885.55, respectively. In contrast, the boundaries between these layers were ambiguous in the corresponding ion distributions for the normal sample (Fig. 3b). In addition, the high spatial resolution is reflected by the observed fibrous myelin extended to the granular layer in the ion distribution of [SHexCer d18:1/24:0 − H]^−^ (Fig. 3c). These results demonstrate that our expansion DESI MSI method achieved single-cell level resolution in mouse brain tissues, enabling the visualization of detailed structures that surpasses the capabilities of conventional DESI MSI. The ability to resolve such fine structural details highlights the significant advancement of our Ex-DESI MSI approach, offering a powerful tool for detailed molecular imaging in complex biological systems.

**Fig. 3:**
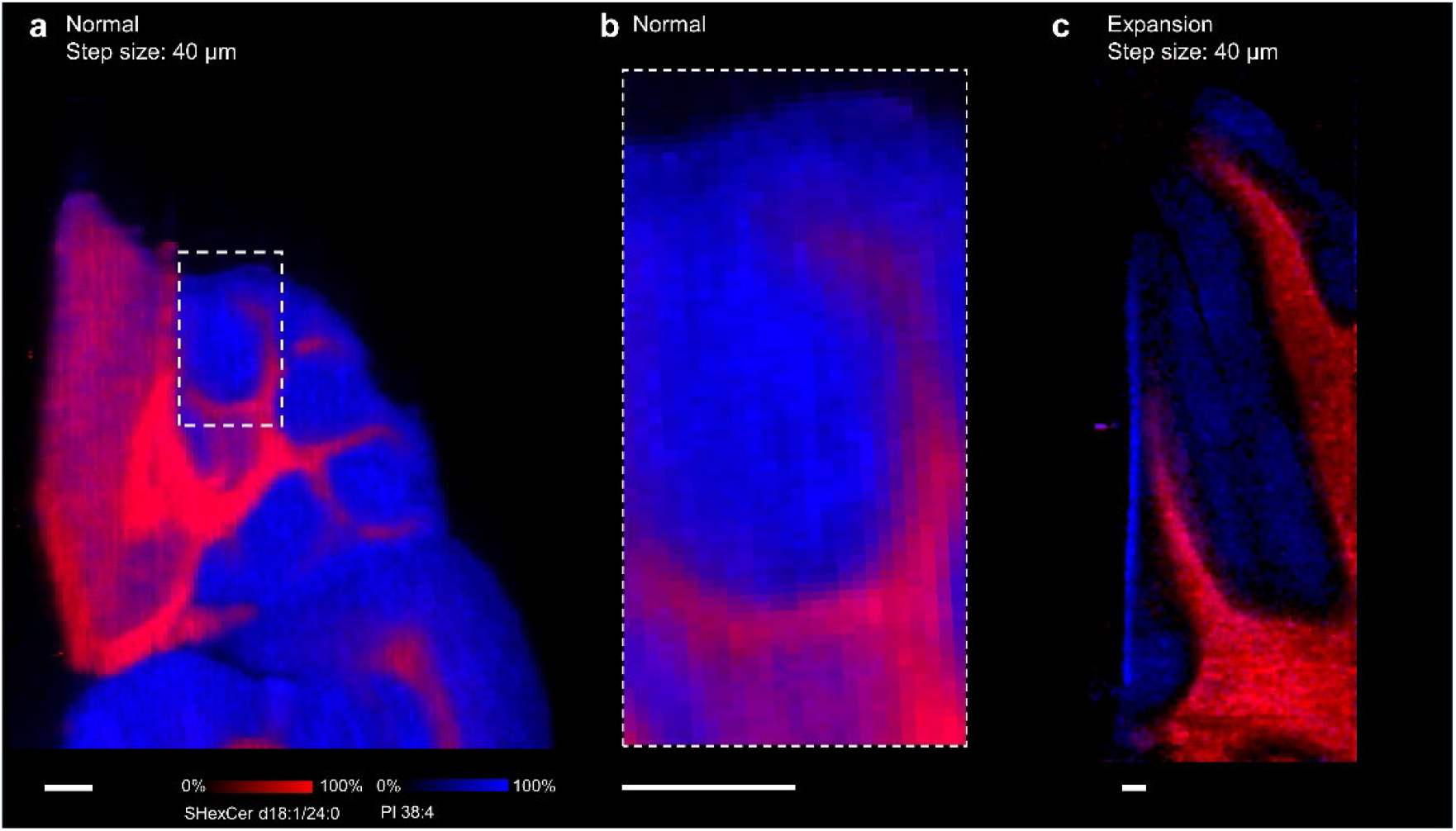
High lateral resolution expansion DESI MSI of mouse cerebellum. Overlay ion images of [SHexCer d18:1/24:0 – H]^−^ (m/z 888.62) and [PI 38:4-H] ^−^ (m/z 885.55) for mouse cerebellum acquired by a) conventional DESI MSI and c) expansion DESI MSI at a step size of 40 μm and line speed of 19.2 μm/s. The ion images for expansion sample were treated with weak denoising. b) expanded view of the region indicated with a white box in a. Scale bar represents 2 mm.

### Lipid assignment for Ex-DESI

An essential aspect of validating any sample expansion technique is ensuring the retention and detectability of molecular species after expansion and sample preparation. To assess the effectiveness of our Ex-DESI MSI method in preserving molecular information, all lipids detected on expanded mouse brain tissues were further annotated by searching LIPID MAPS with a mass tolerance of 5 ppm.. For the mouse brain measured after expansion, abundant lipid species can be observed in the averaged mass spectrum in Fig. 4a from the whole mouse hippocampus region after expansion. The assignment of lipid species is listed in Supplementary Table 1. Phosphatidylserine (PS), phosphatidic acid (PA), and Hexosylceramide (HexCer) count the most in the assigned lipid species for the expansion tissue. In the comparison with the specie assignment for unexpanded sample (Fig. 4b), a significant number loss is observed for Phosphatidylethanolamine (PE) and Phosphatidylinositol (PI) species, possibly due to the fixation process with paraformaldehyde during mouse brain pretreatment. However, an increased assigned number is observed for HexCer and Ceramide (Cer) in the expanded sample (Fig. 4b). This increase may be attributed to changes in the chemical microenvironment during the pretreatment process, as well as to the loss of high-abundance lipids that could suppress the signals of low-abundance species, thereby enhancing their detectability.

**Fig. 4:**
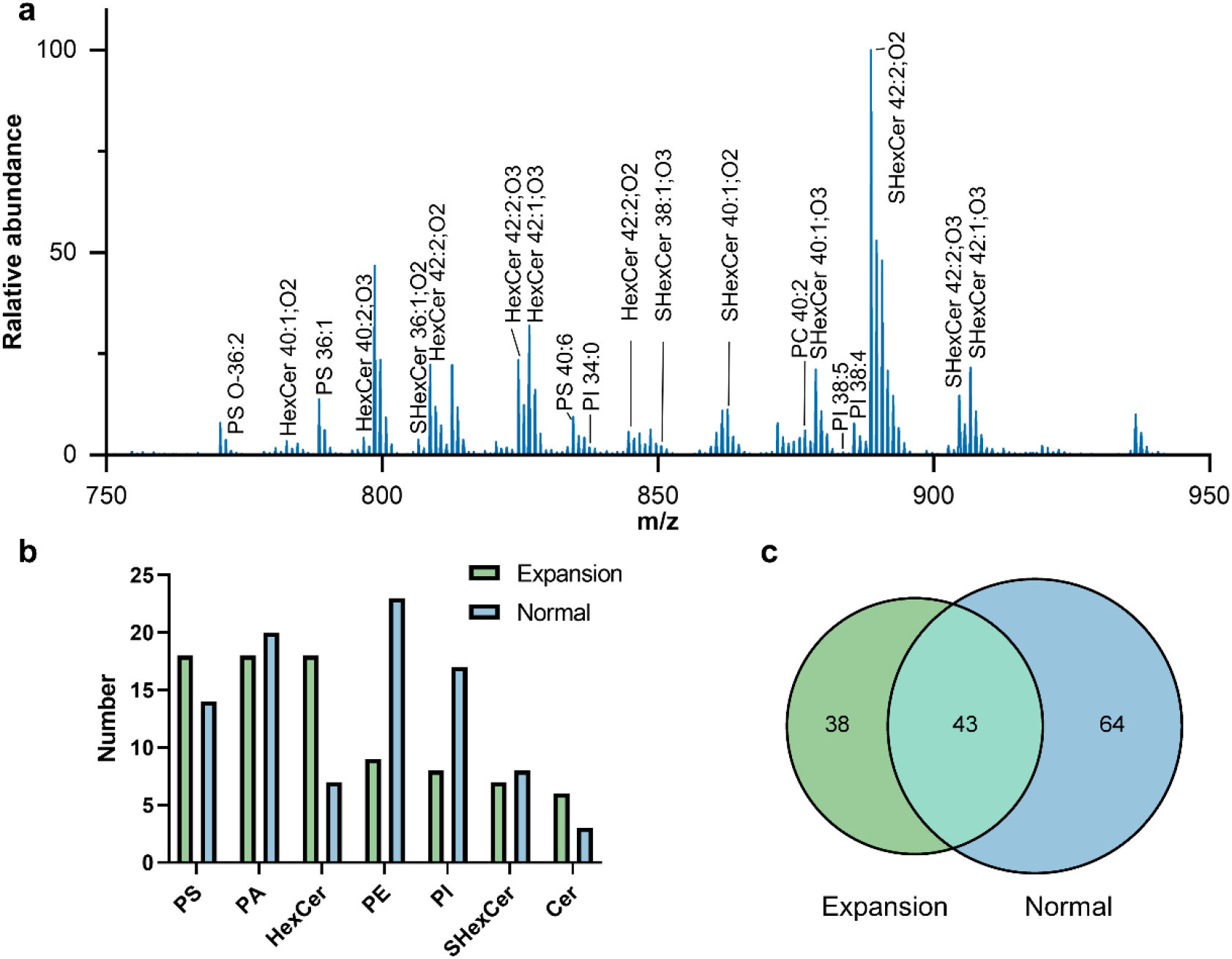
Lipid assignment for expansion and conventional DESI MSI. a) Averaged mass spectrum for expanded mouse hippocampus using a step size of 100 μm in the negative ion mode. b) Number of assigned species for different lipid classes (top 7) from the expanded mouse hippocampus tissue slide and conventional counterpart. Specific assignments are listed in Suppl. Table 1 and 2. c) Comparison of assigned number between expansion and normal sample.

In total, 81 lipid species were assigned for expanded mouse brain tissue, compared to 107 species for conventional brain tissue (Fig. 4c). Despite the decrease in certain lipid classes, the overall lipid retention was high, and many lipid species remained identifiable after tissue expansion. The results demonstrate the high lipid retention efficiency of our Ex-DESI MSI method, even after complex sample pretreatment.

## Conclusion

In summary, this study introduced a cellular-level resolution DESI-MS imaging approach, which provides a more accessible method for analyzing complex biological samples at high spatial resolution, representing a significant improvement compared to traditional DESI MSI techniques. This approach overcomes the longstanding spatial resolution limitations of conventional DESI MSI, enabling the visualization of fine structural details in complex biological tissues, such as mouse brain. Our Ex-DESI MSI not only significantly enhances spatial resolution but also maintains robust molecular detection capabilities.

The integration of high spatial resolution and effective molecular retention underscores the potential of Ex-DESI MSI as a powerful and accessible tool for detailed molecular imaging in various biological applications. By physically enlarging the sample, we bypass the inherent constraints of DESI spray geometry and solvent plume size without requiring modifications for existing mass spectrometers. This advancement represents a significant contribution to the field of mass spectrometry imaging, making high-resolution DESI MSI accessible and reliable for researchers aiming to explore cellular-level structures and functions..

## Methods

### Reagents and materials

| Reagent | Acronym | Vendor | Catalog Number |
| --- | --- | --- | --- |
| Sodium acrylate | SA | Santa Cruz Biotechnology | sc-236893D |
| Sodium acrylate | SA | AK Scientific | R624 |
| N,N-Methylenebisacrylamide | BIS | Sigma Aldrich | M7279 |
| N,N-dimethylacrylamide | DMAA | Sigma Aldrich | 274135 |
| Acrylamide | AA | Sigma Aldrich | A8887 |
| 4-hydroxy-TEMPO | 4HT | Sigma Aldrich | 176141 |
| N,N,N',N' Tetramethylethylenediamine | TEMED | Sigma Aldrich | T9281 |
| Methacrolein |  | Sigma Aldrich | 133035 |
| Phosphate Buffered Saline, 10x Solution, PH 7.4 | PBS | Thermo Fisher Scientific | 70011044 |
| Ammonium persulfate | APS | Sigma Aldrich | A3678 |
| Sodium chloride | NaCl | Sigma Aldrich | S6191 |
| Triton X-100 |  | Sigma Aldrich | T8787 |
| Proteinase K (PCR grade) | ProK | Thermo Fisher Scientific | EO0492 |
| Tris-HCl, 1 M, pH 8 |  | Thermo Fisher Scientific | 15568025 |
| 4% Paraformaldehyde Fix Solution |  | Beyotime Biotechnology | P0099 |
| sucrose |  | Sigma Aldrich | S112228 |
| Dimethyl-formamide | DMF | TIC | D0939 |
| Methanol (HPLC-grade) | MeOH | VWR | BDH20864 |

Superfrost Plus Adhesion Slides was purchased from Epredia (Kalamazoo, MI). 22×22 mm square cover glass was obtained from Corning (Corning, NY). Large plain slides with dimensions of 75 × 52 mm were acquired from Citotest Scientific (Jiangsu, China). Fused silica capillaries were obtained from Molex (Lisle, IL). NanoTight sleeves, PEEK needle port, fingertight fitting, PEEK tee, PEEK unions, and 1/16’’ stainless steel tubing were acquired from IDEX Health & Sciences (Oak Harbor, WA). A 2.5 mL syringe model 1002 was obtained from Hamilton (Reno, NV).

### Animal

C57/BL mice were obtained from the Laboratory Animal Service Centre at the Chinese University of Hong Kong. The mice were maintained under standard conditions, and provided with sterilized water and standard laboratory chow. All animal procedures were conducted with the approval of the Hong Kong Baptist University Committee on the Use of Human and Animal Subjects in Teaching and Research.

### Preparation of tissue slides

Following the sacrifice of the mice, brains were dissected and postfixed overnight at 4 °C in a solution of 4% paraformaldehyde (PFA) in 1x PBS. The organs were then washed in PBS for an additional night at 4 °C. Then the organs were incubated in 15% and subsequently 30% (w/v) sucrose in 1x PBS at 4 °C until they settled at the bottom. The treated tissues were stored at −80 °C until sectioning. 50 μm and 10 μm sections were prepared for expansion and non-expansion experiments, respectively, using a CryoStar Nx70 cryostat (Thermal Fisher Scientific, Walldorf, Germany) set to −20 °C.

### Gelation

A fresh monomer solution was prepared consisting of 340 mg/mL sodium acrylate, 100 mg/mL acrylamide, 0.1 mg/mL N,N’-methylenebisacrylamide, 40 mg/mL N,N’-dimethylacrylamide, and 10 mg/mL sodium chloride in 1x PBS. Prior to gelation, tissue slices were placed into a custom gelling chamber made of two spacers cut from cover glass attached to a microscope slide. Before initiating gelation, 4HT, TEMED, APS, and methacrolein were added to the monomer solution to achieve final concentrations of 0.25% APS (w/v), 0.001% 4HT(w/v), 0.04% TEMED (v/v), and 0.1% (v/v) methacrolein, with TEMED and APS added last to avoid premature gelation. The tissue was immersed in the gelling solution for 30 minutes at 4 °C to allow the monomer solution to penetrate. A glass microscope slide wrapped in parafilm was placed over the gelling chamber, and the assembly was incubated overnight in a humidified container at 37 °C.

### Expansion

The excess gel surrounding the tissue slice was trimmed into an asymmetric shape. The gelation chamber was then placed into a custom-built slide box containing 6 units/mL proteinase K in a digesting solution made of 25 mM EDTA (pH 8), 50 mM Tris-Cl (pH 8), 0.8 M NaCl, and 0.5% Triton-X 100 to perform digestion at 60 °C for 1.5 hours in a humid chamber. The gel was carefully detached and flipped upside down so that the tissue slice faced up. The gel was rinsed three times with 4 °C PBS for 15 minutes each on ice, and then expanded in 4 °C deionized water for 20 minutes, repeated four times on ice, until no further expansion was observed. The gel was rinsed quickly three times with deionized water. After rinsing, the slide box was placed in a drying chamber filled with silica gel desiccant beads under gentle nitrogen gas flow.

### DESI analysis

Expansion imaging utilize DESI MS for spatial profiling of expanded tissue slides. The DESI probe was home-built comprising of a 1/16’’ stainless-steel sheath gas tube and an inter fused silica capillary of 150/20 μm (O.D./I.D.) to deliver solvent (Fig. 1b). The capillary was etched by hydrofluoric acid (HF) following the steps reported^15^. In briefly, the end of the capillary was burned and insert into a 49% aqueous HF solution. Water is delivered through the capillary at a flow rate of 0.1 μL/min to prevent etchant from entering the interior of the capillary. A tapered capillary tip is created once the etch process is halted by removing the silica touching the solution surface. The solvent capillary extended through a high pressure PEEK Tee (0.020” thru hole) assembly with Fingertight Fittings for 1/16” OD Tubing and NanoTight Tubing Sleeves (1/16’’ tubing O.D. with 0.007’’ I.D.). Sheath gas was induced through the middle hole of PEEK Tee. To reduce back pressure, the capillary of 20 μm I.D. was only used in the DESI probe and connected to a fused silica capillary of 360/100 μm (O.D./I.D.) using a stainless union where also the high voltage is applied. A PEEK needle port was used to connected the syringe and capillary through a stainless union.

A solvent mixture of 1:4(v/v) dimethylformamide : methanal at a flow rate of 1.5 μL/min was used. Sheath gas was set to 250 psi. The DESI probe was positioned 6.5 mm away from the extended ion transfer tube (ITT) and at the same height as the top of the ITT at an angle of 70 °.

Mass spectrum was acquired at a negative ion mode by Q-Exactive mass spectrometer (Themo Fisher). Spray voltage was set to −4000V. Mass resolution was 3,5000 at m/z 200 with scanning mass range of 600-1500. Capillary temperature was 400 ^°^C and S-lens RF voltage is −100V. The ion injection time was 200 ms. A custom-built moving system was used to control the movement of sample stage.

### DESI-MSI data analysis

A complete MSI file was produced by one .raw file. Firstly, .raw file acquired by mass spectrometer was converted into .mzML file using MSConvert (ProteoWizard), and then transfer into .imzML file using MT imzML Converter (ng) 1.4.1 (MassTech). One pixel was produced from two spectra by picking up max TIC spectrum. The pixel width was calculated from the lateral speed of the moving stage and the cycle time of MS while pixel height was determined by the vertical step size. imzML file was visualized by SCILS Lab MVS (2024a Premium 3D). All ion images were was normalized to the total ion current (TIC).

### Lipid assignment

Identified ion peaks in the averaged mass spectra from the whole measured tissue slides were used for lipid assignment. LIPID MAPS® Structure Database (LMSD) from LIPID MAP was used for specie assignment by standard library search. Ion adduct forms of [M-H]^−^ and [M+Cl]^−^ were considered in this study. Artificial screening was conducted to exclude fatty acyl chains with odd length, non-mammalian and unreported species.

## Acknowledgments

This work was supported by General Research Fund (12302122) of the Research Grants Council, Hong Kong Special Administrative Region, SKLEBA Research Grant (SKLP_2021_P04), and a Start-up Grant from Hong Kong Baptist University.

A US provisional patent application has been filed covering the technology/methodology described in this manuscript.

