## Supplemental Information for "Single-Cell Resolution DESI Mass Spectrometry Imaging through 10-Fold Sample Expansion"

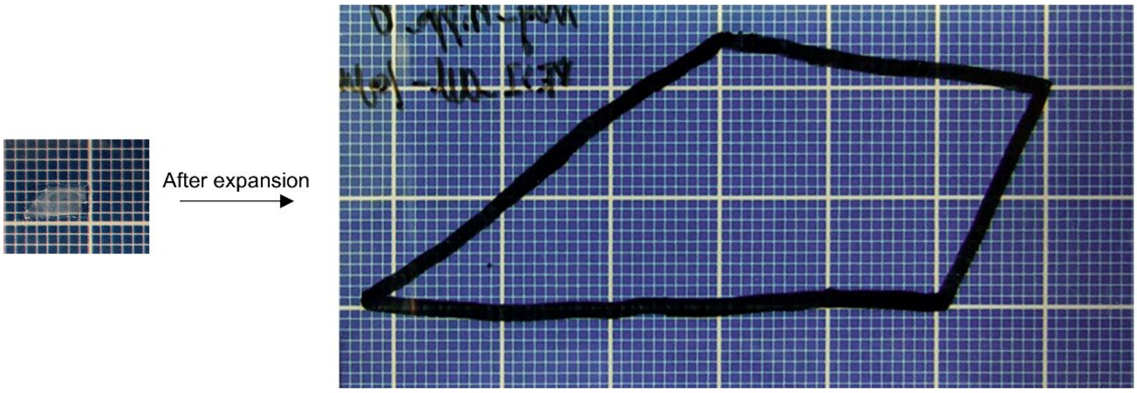


**Supplementary Fig. 1**: Tissue section before and after expansion.

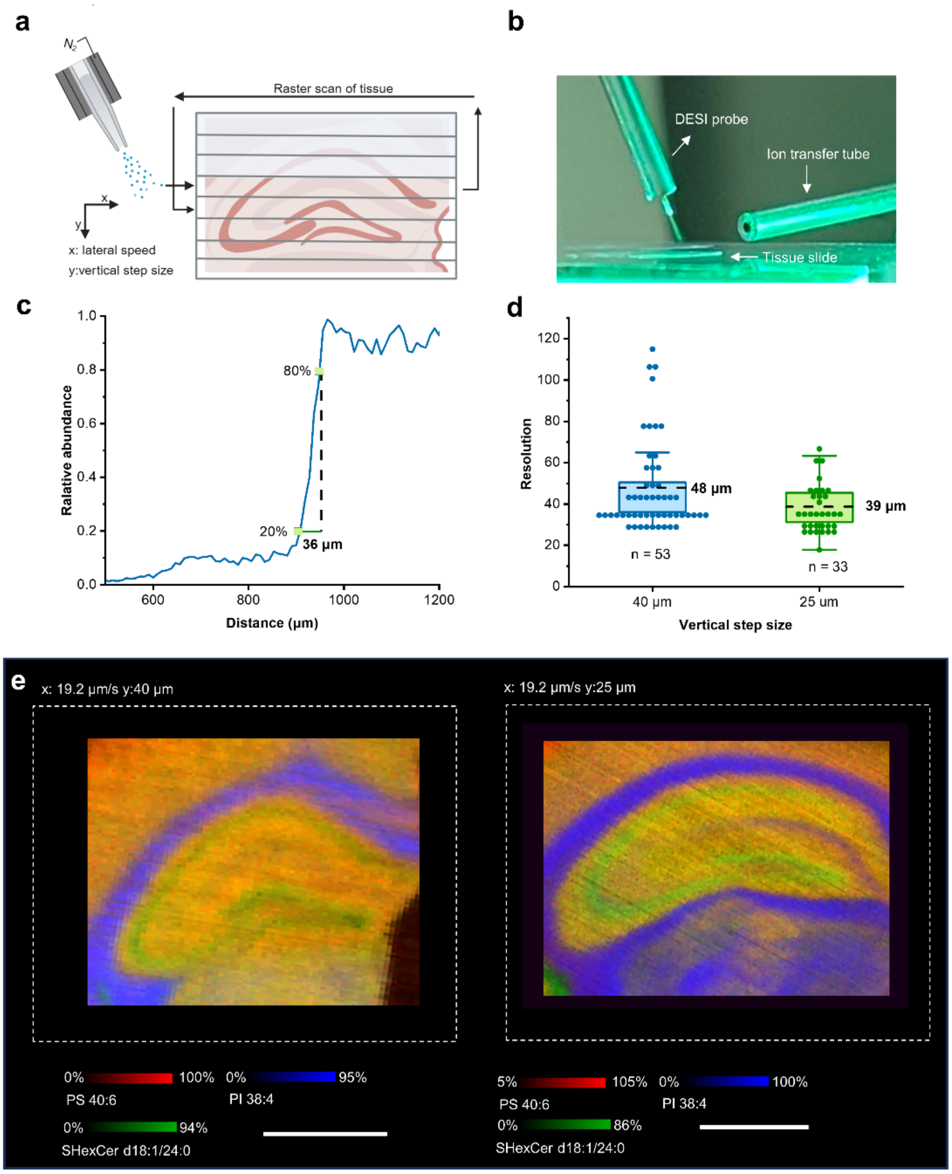


**Supplementary Fig. 2**: **Determination of the lateral resolution for DESI MSI.** a) Schematic of the scanning mode of DESI MSI shows that continuous spray scans across the tissue section row by row to finish the imaging process. b) photograph of the DESI imaging process. c) Demonstration of the 20-80% rule for lateral resolution measurement using the extracted ion chromatogram of PS 40:5 from one row of the mouse hippocampus tissue. d) Lateral resolution for mouse brain tissues using the step size of 40 μm and 25 μm. Dashed lines and labels shows the mean values. The box contains the 25th-75th percentile. 53 rows were counted for 40 μm step size, while 33 rows for 25 μm step size. e) overlay ion images of PS 40:6 (m/z 834.53), PI

38:4 (m/z 885.55), and SHexCer d18:1/24:0 (m/z 888.62) obtained from normal mouse hippocampus using step size of 40 µm and 25 µm. Scale bar represents 1mm.

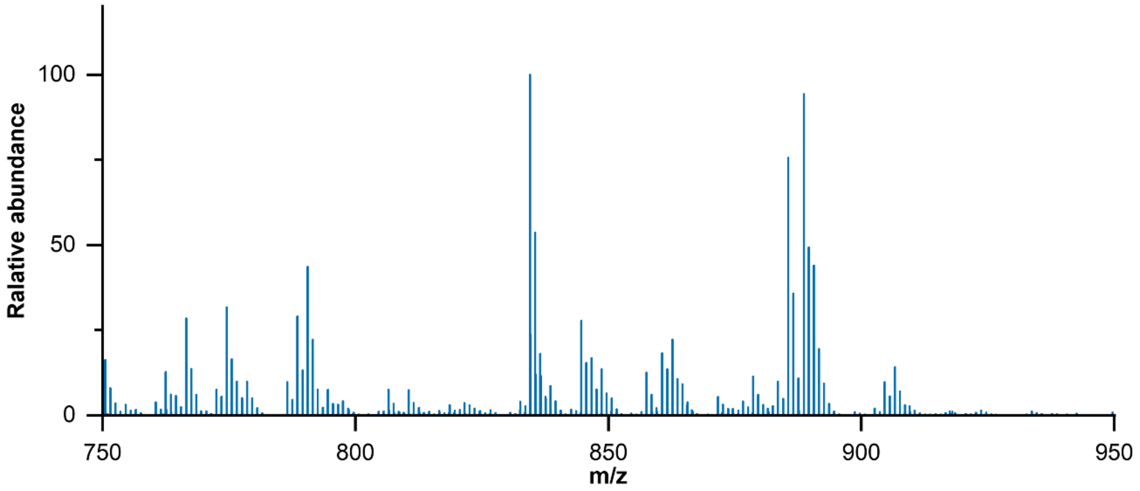


**Supplementary Fig. 3**: Averaged mass spectrum for normal mouse hippocampus using a step size of 100 µm in the negative ion mode.

Supplementary Table 1. List of lipid assignments within 5 ppm mass error for the ion species detected in expanded mouse brain slice under negative mode. Step size was 100 µm.

| Theoretical m/z | Experimental m/z | Error (ppm) | Identity | Chemical Formula | Ion Type |
|---|---|---|---|---|---|
| 608.5629 | 608.5623 | 1.0 | Cer 38:1;O3 | C38H75NO4 | $[M-H]^-$ |
| 618.584 | 618.5831 | 1.5 | Cer 40:2;O2 | C40H77NO3 | $[M-H]^-$ |
| 620.5996 | 620.5987 | 1.5 | Cer 40:1;O2 | C40H79NO3 | $[M-H]^-$ |
| 636.5943 | 636.5936 | 1.1 | Cer 40:1;O3 | C40H79NO4 | $[M-H]^-$ |
| 646.6153 | 646.6144 | 1.4 | Cer 42:2;O2 | C42H81NO3 | $[M-H]^-$ |
| 647.4666 | 647.4657 | 1.4 | PA 32:0 | C35H69O8P | $[M-H]^-$ |
| 669.45 | 669.4501 | 0.1 | PA 34:3 | C37H67O8P | $[M-H]^-$ |
| 673.4825 | 673.4814 | 1.8 | PA 34:1 | C37H71O8P | $[M-H]^-$ |
| 682.5889 | 682.5911 | 3.1 | Cer 42:2;O2 | C42H81NO3 | $[M+Cl]^-$ |
| 683.504 | 683.5021 | 2.8 | PA O-36:3 | C39H73O7P | $[M-H]^-$ |
| 685.5193 | 685.5178 | 2.2 | PA O-36:2 | C39H75O7P | $[M-H]^-$ |
| 695.4652 | 695.4657 | 0.9 | PA 36:4 | C39H69O8P | $[M-H]^-$ |
| 699.4975 | 699.497 | 0.7 | PA 36:2 | C39H73O8P | $[M-H]^-$ |
| 701.5131 | 701.5127 | 0.6 | PA 36:1 | C39H75O8P | $[M-H]^-$ |
| 707.5056 | 707.5021 | 4.9 | PA O-38:5 | C41H73O7P | $[M-H]^-$ |
| 711.5362 | 711.5334 | 3.8 | PA O-38:3 | C41H77O7P | $[M-H]^-$ |
| 716.5254 | 716.5236 | 2.5 | PE 34:1 | C39H76NO8P | $[M-H]^-$ |
| 717.4467 | 717.4501 | 4.7 | PA 38:7 | C41H67O8P | $[M-H]^-$ |
| 718.5392 | 718.5392 | 0.0 | PE 34:0 | C39H78NO8P | $[M-H]^-$ |
| 720.4958 | 720.4974 | 2.2 | PE O-36:6 | C41H72NO7P | $[M-H]^-$ |
| 723.4972 | 723.497 | 0.3 | PA 38:4 | C41H73O8P | $[M-H]^-$ |
| 727.5298 | 727.5283 | 2.1 | PA 38:2 | C41H77O8P | $[M-H]^-$ |
| 734.4949 | 734.4978 | 3.8 | PS 32:0 | C38H74NO10P | $[M-H]^-$ |
| 736.4902 | 736.4923 | 2.9 | PE 36:5 | C41H72NO8P | $[M-H]^-$ |
| 738.5046 | 738.5079 | 4.5 | PE 36:4 | C41H74NO8P | $[M-H]^-$ |
| 740.5217 | 740.5236 | 2.6 | PE 36:3 | C41H76NO8P | $[M-H]^-$ |
| 744.5547 | 744.5549 | 0.3 | PE 36:1 | C41H80NO8P | $[M-H]^-$ |
| 745.4796 | 745.4814 | 2.4 | PA 40:7 | C43H71O8P | $[M-H]^-$ |
| 747.5073 | 747.5101 | 3.7 | PA O-38:3 | C41H77O7P | $[M+Cl]^-$ |
| 751.5304 | 751.5283 | 2.8 | PA 40:4 | C43H77O8P | $[M-H]^-$ |

| | | | | | |
|---|---|---|---|---|---|
| 752.6013 | 752.6046 | 4.4 | HexCer 38:2;O2 | C44H83NO8 | $[M-H]^-$ |
| 753.5476 | 753.544 | 4.9 | PA 40:3 | C43H79O8P | $[M-H]^-$ |
| 754.6217 | 754.6202 | 2.0 | HexCer 38:1;O2 | C44H85NO8 | $[M-H]^-$ |
| 756.4941 | 756.4952 | 1.5 | PS O-32:0 | C38H76NO9P | $[M+Cl]^-$ |
| 756.6328 | 756.6359 | 4.1 | HexCer 38:0;O2 | C44H87NO8 | $[M-H]^-$ |
| 762.5067 | 762.5079 | 1.6 | PE 38:6 | C43H74NO8P | $[M-H]^-$ |
| 762.5657 | 762.5656 | 0.0 | HexCer 36:1;O2 | C42H81NO8 | $[M+Cl]^-$ |
| 766.4419 | 766.4431 | 1.6 | PS 32:2 | C38H70NO10P | $[M+Cl]^-$ |
| 766.5409 | 766.5392 | 2.2 | PE 38:4 | C43H78NO8P | $[M-H]^-$ |
| 769.5005 | 769.5025 | 2.6 | PG 36:4 | C42H75O10P | $[M-H]^-$ |
| 769.6079 | 769.6117 | 4.9 | PA O-42:2 | C45H87O7P | $[M-H]^-$ |
| 772.5514 | 772.5498 | 2.1 | PS O-36:2 | C42H80NO9P | $[M-H]^-$ |
| 778.477 | 778.4795 | 3.3 | PS O-34:3 | C40H74NO9P | $[M+Cl]^-$ |
| 780.6366 | 780.6359 | 0.9 | HexCer 40:2;O2 | C46H87NO8 | $[M-H]^-$ |
| 782.6523 | 782.6515 | 0.9 | HexCer 40:1;O2 | C46H89NO8 | $[M-H]^-$ |
| 786.5307 | 786.5291 | 2.0 | PS 36:2 | C42H78NO10P | $[M-H]^-$ |
| 788.5456 | 788.5447 | 1.1 | PS 36:1 | C42H80NO10P | $[M-H]^-$ |
| 793.5241 | 793.5236 | 0.5 | PI O-32:1 | C41H79O12P | $[M-H]^-$ |
| 794.536 | 794.5342 | 2.4 | PS O-38:5 | C44H78NO9P | $[M-H]^-$ |
| 795.5378 | 795.5393 | 1.9 | PI O-32:0 | C41H81O12P | $[M-H]^-$ |
| 796.6304 | 796.6308 | 0.6 | HexCer 40:2;O3 | C46H87NO9 | $[M-H]^-$ |
| 800.5832 | 800.5811 | 2.6 | PS O-38:2 | C44H84NO9P | $[M-H]^-$ |
| 801.5783 | 801.5805 | 2.7 | TG 46:6 | C49H82O6 | $[M+Cl]^-$ |
| 806.5417 | 806.5458 | 5.0 | SHexCer 36:1;O2 | C42H81NO11S | $[M-H]^-$ |
| 808.6682 | 808.6672 | 1.2 | HexCer 42:2;O2 | C48H91NO8 | $[M-H]^-$ |
| 809.5157 | 809.5186 | 3.6 | PI 32:0 | C41H79O13P | $[M-H]^-$ |
| 810.5288 | 810.5291 | 0.2 | PS 38:4 | C44H78NO10P | $[M-H]^-$ |
| 810.6837 | 810.6828 | 1.1 | HexCer 42:1;O2 | C48H93NO8 | $[M-H]^-$ |
| 814.5617 | 814.5604 | 1.6 | PS 38:2 | C44H82NO10P | $[M-H]^-$ |
| 816.5768 | 816.576 | 0.9 | PS 38:1 | C44H84NO10P | $[M-H]^-$ |
| 818.5355 | 818.5342 | 1.6 | PS O-40:7 | C46H78NO9P | $[M-H]^-$ |
| 818.6256 | 818.6282 | 3.2 | HexCer 40:1;O2 | C46H89NO8 | $[M+Cl]^-$ |

| | | | | | |
|---|---|---|---|---|---|
| 820.5475 | 820.5498 | 2.8 | PS O-40:6 | C46H80NO9P | [M-H]- |
| 824.6627 | 824.6621 | 0.7 | HexCer 42:2;O3 | C48H91NO9 | [M-H]- |
| 826.6776 | 826.6778 | 0.2 | HexCer 42:1;O3 | C48H93NO9 | [M-H]- |
| 832.6111 | 832.6075 | 4.3 | HexCer 40:2;O3 | C46H87NO9 | [M+Cl]- |
| 834.5299 | 834.5291 | 1.1 | PS 40:6 | C46H78NO10P | [M-H]- |
| 835.531 | 835.5342 | 3.8 | PI 34:1 | C43H81O13P | [M-H]- |
| 837.5458 | 837.5499 | 4.9 | PI 34:0 | C43H83O13P | [M-H]- |
| 838.5589 | 838.5604 | 1.8 | PS 40:4 | C46H82NO10P | [M-H]- |
| 839.5613 | 839.5574 | 4.5 | PG 38:1 | C44H85O10P | [M+Cl]- |
| 844.6456 | 844.6439 | 2.0 | HexCer 42:2;O2 | C48H91NO8 | [M+Cl]- |
| 850.5717 | 850.572 | 0.4 | SHexCer 38:1;O3 | C44H85NO12S | [M-H]- |
| 856.5111 | 856.5134 | 2.7 | PS 42:9 | C48H76NO10P | [M-H]- |
| 857.5158 | 857.5186 | 3.3 | PI 36:4 | C45H79O13P | [M-H]- |
| 862.6127 | 862.6084 | 5.0 | SHexCer 40:1;O2 | C46H89NO11S | [M-H]- |
| 868.6247 | 868.6204 | 5.0 | PS O-40:0 | C46H92NO9P | [M+Cl]- |
| 876.6238 | 876.6255 | 1.9 | PC 40:2 | C48H92NO8P | [M+Cl]- |
| 878.6036 | 878.6033 | 0.3 | SHexCer 40:1;O3 | C46H89NO12S | [M-H]- |
| 883.5313 | 883.5342 | 3.3 | PI 38:5 | C47H81O13P | [M-H]- |
| 885.5496 | 885.5499 | 0.3 | PI 38:4 | C47H83O13P | [M-H]- |
| 888.6239 | 888.624 | 0.1 | SHexCer 42:2;O2 | C48H91NO11S | [M-H]- |
| 904.6186 | 904.6189 | 0.3 | SHexCer 42:2;O3 | C48H91NO12S | [M-H]- |
| 906.6343 | 906.6346 | 0.3 | SHexCer 42:1;O3 | C48H93NO12S | [M-H]- |
| 1380.002 | 1379.996 | 3.9 | CL 66:0 | C75H146O17P2 | [M-H]- |
| 1437.961 | 1437.957 | 2.6 | CL 68:3 | C77H144O17P2 | [M+Cl]- |

Supplementary Table 2. List of lipid assignments within 5 ppm mass error for the ion species detected in normal mouse brain slice under negative mode. Step size was 100 μm.

| Theoretical m/z | Experimental m/z | Error (ppm) | Identity | Chemical Formula | Ion Type |
|---|---|---|---|---|---|
| 618.5843 | 618.5831 | 1.9 | Cer 40:2;O2 | C40H77NO3 | $[M-H]^-$ |
| 620.5989 | 620.5987 | 0.3 | Cer 40:1;O2 | C40H79NO3 | $[M-H]^-$ |
| 645.4501 | 645.4501 | 0.0 | PA 32:1 | C35H67O8P | $[M-H]^-$ |
| 646.6157 | 646.6144 | 2.0 | Cer 42:2;O2 | C42H81NO3 | $[M-H]^-$ |
| 647.3516 | 647.3485 | 4.8 | PA 30:4 | C33H57O8P | $[M+Cl]^-$ |
| 647.4668 | 647.4657 | 1.5 | PA 32:0 | C35H69O8P | $[M-H]^-$ |
| 657.4878 | 657.4865 | 2.0 | PA O-34:2 | C37H71O7P | $[M-H]^-$ |
| 659.5026 | 659.5021 | 0.6 | PA O-34:1 | C37H73O7P | $[M-H]^-$ |
| 669.4491 | 669.4501 | 1.5 | PA 34:3 | C37H67O8P | $[M-H]^-$ |
| 671.466 | 671.4657 | 0.3 | PA 34:2 | C37H69O8P | $[M-H]^-$ |
| 672.4971 | 672.4974 | 0.4 | PE O-32:2 | C37H72NO7P | $[M-H]^-$ |
| 673.4825 | 673.4814 | 1.6 | PA 34:1 | C37H71O8P | $[M-H]^-$ |
| 679.4135 | 679.4111 | 3.4 | PA 32:2 | C35H65O8P | $[M+Cl]^-$ |
| 683.503 | 683.5021 | 1.3 | PA O-36:3 | C39H73O7P | $[M-H]^-$ |
| 685.5186 | 685.5178 | 1.3 | PA O-36:2 | C39H75O7P | $[M-H]^-$ |
| 690.5074 | 690.5079 | 0.9 | PE 32:0 | C37H74NO8P | $[M-H]^-$ |
| 695.4658 | 695.4657 | 0.0 | PA 36:4 | C39H69O8P | $[M-H]^-$ |
| 697.4801 | 697.4814 | 1.9 | PA 36:3 | C39H71O8P | $[M-H]^-$ |
| 698.5143 | 698.513 | 1.9 | PE O-34:3 | C39H74NO7P | $[M-H]^-$ |
| 699.4983 | 699.497 | 1.9 | PA 36:2 | C39H73O8P | $[M-H]^-$ |
| 707.5024 | 707.5021 | 0.4 | PA O-38:5 | C41H73O7P | $[M-H]^-$ |
| 711.5349 | 711.5334 | 2.0 | PA O-38:3 | C41H77O7P | $[M-H]^-$ |
| 713.5521 | 713.5491 | 4.2 | PA O-38:2 | C41H79O7P | $[M-H]^-$ |
| 714.5078 | 714.5079 | 0.1 | PE 34:2 | C39H74NO8P | $[M-H]^-$ |
| 716.5247 | 716.5236 | 1.7 | PE 34:1 | C39H76NO8P | $[M-H]^-$ |
| 718.5398 | 718.5392 | 0.8 | PE 34:0 | C39H78NO8P | $[M-H]^-$ |
| 722.5149 | 722.513 | 2.6 | PE O-36:5 | C41H74NO7P | $[M-H]^-$ |
| 726.5429 | 726.5443 | 2.1 | PE O-36:3 | C41H78NO7P | $[M-H]^-$ |
| 735.4746 | 735.4737 | 1.2 | PA 36:2 | C39H73O8P | $[M+Cl]^-$ |
| 735.5364 | 735.5334 | 4.1 | PA O-40:5 | C43H77O7P | $[M-H]^-$ |

| | | | | | |
|---|---|---|---|---|---|
| 738.5083 | 738.5079 | 0.4 | PE 36:4 | C41H74NO8P | $[M-H]^-$ |
| 744.556 | 744.5549 | 1.5 | PE 36:1 | C41H80NO8P | $[M-H]^-$ |
| 746.5137 | 746.513 | 0.9 | PE O-38:7 | C43H74NO7P | $[M-H]^-$ |
| 747.5179 | 747.5182 | 0.3 | PG 34:1 | C40H77O10P | $[M-H]^-$ |
| 748.5272 | 748.5287 | 2.0 | PE O-38:6 | C43H76NO7P | $[M-H]^-$ |
| 750.5463 | 750.5443 | 2.7 | PE O-38:5 | C43H78NO7P | $[M-H]^-$ |
| 754.5756 | 754.5756 | 0.1 | PE O-38:3 | C43H82NO7P | $[M-H]^-$ |
| 760.5161 | 760.5134 | 3.6 | PS 34:1 | C40H76NO10P | $[M-H]^-$ |
| 762.5089 | 762.5079 | 1.3 | PE 38:6 | C43H74NO8P | $[M-H]^-$ |
| 762.5641 | 762.5656 | 2.0 | HexCer 36:1;O2 | C42H81NO8 | $[M+Cl]^-$ |
| 764.5206 | 764.5236 | 3.9 | PE 38:5 | C43H76NO8P | $[M-H]^-$ |
| 766.5403 | 766.5392 | 1.3 | PE 38:4 | C43H78NO8P | $[M-H]^-$ |
| 769.5004 | 769.5025 | 2.7 | PG 36:4 | C42H75O10P | $[M-H]^-$ |
| 772.5292 | 772.5287 | 0.8 | PE O-40:8 | C45H76NO7P | $[M-H]^-$ |
| 772.5893 | 772.5862 | 4.0 | PE 38:1 | C43H84NO8P | $[M-H]^-$ |
| 773.5332 | 773.5338 | 0.9 | PG 36:2 | C42H79O10P | $[M-H]^-$ |
| 774.5457 | 774.5443 | 1.7 | PE O-40:7 | C45H78NO7P | $[M-H]^-$ |
| 778.5745 | 778.5756 | 1.4 | PE O-40:5 | C45H82NO7P | $[M-H]^-$ |
| 785.6083 | 785.6066 | 2.2 | PA 42:1 | C45H87O8P | $[M-H]^-$ |
| 786.5323 | 786.5291 | 4.2 | PS 36:2 | C42H78NO10P | $[M-H]^-$ |
| 788.545 | 788.5447 | 0.4 | PS 36:1 | C42H80NO10P | $[M-H]^-$ |
| 790.5415 | 790.5392 | 2.9 | PE 40:6 | C45H78NO8P | $[M-H]^-$ |
| 794.5695 | 794.5705 | 1.3 | PE 40:4 | C45H82NO8P | $[M-H]^-$ |
| 796.6308 | 796.6308 | 0.0 | HexCer 40:2;O3 | C46H87NO9 | $[M-H]^-$ |
| 797.5371 | 797.5338 | 4.1 | PG 38:4 | C44H79O10P | $[M-H]^-$ |
| 806.5456 | 806.5458 | 0.1 | SHexCer 36:1;O2 | C42H81NO11S | $[M-H]^-$ |
| 807.557 | 807.5546 | 3.0 | PG O-40:6 | C46H81O9P | $[M-H]^-$ |
| 808.6686 | 808.6672 | 1.7 | HexCer 42:2;O2 | C48H91NO8 | $[M-H]^-$ |
| 809.5068 | 809.5105 | 4.6 | PG 36:2 | C42H79O10P | $[M+Cl]^-$ |
| 810.5282 | 810.5291 | 1.1 | PS 38:4 | C44H78NO10P | $[M-H]^-$ |
| 810.6833 | 810.6828 | 0.5 | HexCer 42:1;O2 | C48H93NO8 | $[M-H]^-$ |
| 814.5604 | 814.5604 | 0.0 | PS 38:2 | C44H82NO10P | $[M-H]^-$ |
| 816.5745 | 816.576 | 1.8 | PS 38:1 | C44H84NO10P | $[M-H]^-$ |

| | | | | | |
|---|---|---|---|---|---|
| 819.5291 | 819.5312 | 2.6 | PG O-38:4 | C44H81O9P | [M+Cl]- |
| 821.5547 | 821.5549 | 0.2 | PI O-34:1 | C43H83O12P | [M-H]- |
| 824.6633 | 824.6621 | 1.5 | HexCer 36:1;O | C48H91NO9 | [M-H]- |
| 826.6789 | 826.6778 | 1.3 | HexCer 42:1;O3 | C48H93NO9 | [M-H]- |
| 832.5124 | 832.5134 | 1.2 | PS 40:7 | C46H76NO10P | [M-H]- |
| 832.6091 | 832.6075 | 1.9 | HexCer 40:2;O3 | C46H87NO9 | [M+Cl]- |
| 834.5303 | 834.5291 | 1.6 | PS 40:6 | C46H78NO10P | [M-H]- |
| 834.6178 | 834.6149 | 3.5 | PC O-38:2 | C46H90NO7P | [M+Cl]- |
| 838.5577 | 838.5604 | 3.2 | PS 40:4 | C46H82NO10P | [M-H]- |
| 842.5956 | 842.5917 | 4.7 | PS 40:2 | C46H86NO10P | [M-H]- |
| 849.5877 | 849.5862 | 1.6 | PI O-36:1 | C45H87O12P | [M-H]- |
| 850.5742 | 850.572 | 2.6 | SHexCer 38:1;O3 | C44H85NO12S | [M-H]- |
| 852.5562 | 852.5527 | 4.1 | PS 38:1 | C44H84NO10P | [M+Cl]- |
| 854.5814 | 854.5836 | 2.6 | PC O-40:6 | C48H86NO7P | [M+Cl]- |
| 857.5164 | 857.5186 | 2.6 | PI 36:4 | C45H79O13P | [M-H]- |
| 861.5495 | 861.5499 | 0.3 | PI 36:2 | C45H83O13P | [M-H]- |
| 862.6112 | 862.6084 | 3.2 | SHexCer 40:1;O2 | C46H89NO11S | [M-H]- |
| 865.5048 | 865.5025 | 2.7 | PG 44:12 | C50H75O10P | [M-H]- |
| 868.6208 | 868.6204 | 0.5 | PS O-40:0 | C46H92NO9P | [M+Cl]- |
| 874.6084 | 874.6098 | 1.6 | PC 40:3 | C48H90NO8P | [M+Cl]- |
| 876.6248 | 876.6255 | 0.8 | PC 40:2 | C48H92NO8P | [M+Cl]- |
| 877.6183 | 877.6175 | 0.8 | PI O-38:1 | C47H91O12P | [M-H]- |
| 878.6048 | 878.6033 | 1.7 | SHexCer 40:1;O3 | C46H89NO12S | [M-H]- |
| 881.5204 | 881.5186 | 2.0 | PI 38:6 | C47H79O13P | [M-H]- |
| 882.5268 | 882.5291 | 2.6 | PS 44:10 | C50H78NO10P | [M-H]- |
| 883.5316 | 883.5342 | 2.9 | PI 38:5 | C47H81O13P | [M-H]- |
| 885.5509 | 885.5499 | 1.2 | PI 38:4 | C47H83O13P | [M-H]- |
| 888.6256 | 888.624 | 1.8 | SHexCer 42:2;O2 | C48H91NO11S | [M-H]- |
| 890.6366 | 890.6397 | 3.5 | SHexCer 42:1;O2 | C48H93NO11S | [M-H]- |
| 903.4996 | 903.5029 | 3.7 | PI 40:9 | C49H77O13P | [M-H]- |
| 904.62 | 904.6189 | 1.2 | SHexCer 42:2;O3 | C48H91NO12S | [M-H]- |
| 905.5188 | 905.5186 | 0.2 | PI 40:8 | C49H79O13P | [M-H]- |

| | | | | | |
|---|---|---|---|---|---|
| 906.6361 | 906.6346 | 1.7 | SHexCer 42:1;O3 | C48H93NO12S | $[M-H]^-$ |
| 907.5384 | 907.5342 | 4.6 | PI 40:7 | C49H81O13P | $[M-H]^-$ |
| 909.5474 | 909.5499 | 2.7 | PI 40:6 | C49H83O13P | $[M-H]^-$ |
| 914.4711 | 914.4744 | 3.7 | PS 44:12 | C50H74NO10P | $[M+Cl]^-$ |
| 933.6831 | 933.6801 | 3.1 | PI O-42:1 | C51H99O12P | $[M-H]^-$ |
| 935.6937 | 935.6958 | 2.2 | PI O-42:0 | C51H101O12P | $[M-H]^-$ |
| 949.6795 | 949.6751 | 4.6 | PI 42:0 | C51H99O13P | $[M-H]^-$ |
| 953.5183 | 953.5186 | 0.3 | PI 44:12 | C53H79O13P | $[M-H]^-$ |
| 955.6094 | 955.6048 | 4.8 | PI 40:1 | C49H93O13P | $[M+Cl]^-$ |
| 1017.767 | 1017.768 | 1.0 | TG 62:10 | C65H106O6 | $[M+Cl]^-$ |
| 1037.741 | 1037.737 | 3.8 | TG 64:14 | C67H102O6 | $[M+Cl]^-$ |
| 1045.799 | 1045.8 | 1.1 | TG 64:10 | C67H110O6 | $[M+Cl]^-$ |
| 1073.83 | 1073.831 | 1.2 | TG 66:10 | C69H114O6 | $[M+Cl]^-$ |
| 1423.959 | 1423.965 | 4.4 | CL 70:6 | C79H142O17P2 | $[M-H]^-$ |
| 1473.974 | 1473.981 | 4.4 | CL 74:9 | C83H144O17P2 | $[M-H]^-$ |
| 1475.999 | 1475.996 | 1.6 | CL 74:8 | C83H146O17P2 | $[M-H]^-$ |
